# Survey of benzimidazole resistance in ascarid parasites of poultry

**DOI:** 10.1101/2025.08.12.669876

**Authors:** JB Collins, Rachel Choo, Amanda O. Shaver, Etta S. Schaye, Tom Volpe, Lenny Nunn, Megan E. Lighty, Kayla R. Niel, Emily M. Frye, Mostafa Zamanian, Erik C. Andersen

## Abstract

Ascarid parasites, such as *Ascaridia galli* and *Heterakis gallinarum*, are nearly ubiquitous in poultry and can cause serious production losses. *H. gallinarum* is of particular concern because of its role as a vector for the protozoan *Histomonas meleagridis*, the cause of blackhead disease. Currently, only the benzimidazole anthelmintic, fenbendazole (FBZ), is approved for use in poultry, and recently, FBZ resistance has been discovered and validated in populations of the turkey ascarid *Ascaridia dissimilis* and in ascarid of gallinaceous birds *H. gallinarum*. Here, we further explore the prevalence of resistance in poultry ascarids by testing FBZ efficacy against thirteen isolates of *A. galli* and eight isolates of *H. gallinarum*. Isolates were used to infect day-old naive chickens. Four weeks after infection, animals to be treated received the label-recommended dosage of FBZ (SafeGuard Aquasol) for five days, per the manufacturer’s directions. One week after the fifth day of treatment, animals were euthanized and parasite burdens were counted to determine treatment efficacy between the untreated and treated groups. Resistance was identified and validated in a single isolate of *A. galli*, marking the first confirmed case in the species. All isolates of *H. gallinarum* were found to be resistant. The emergence of resistance in *A. galli* and the high prevalence of resistance in *H. gallinarum* highlight the growing concern of resistance in parasites of poultry. Without approved alternative treatments, the detrimental effects of infections cannot be mitigated in resistant populations, significantly impacting profit margins. Diagnostics that enable broader surveys are necessary to understand the full scope of the problem. However, we show that resistance is present across production species and should act as an impetus for the discovery of new treatments and the adoption of new management strategies.

## INTRODUCTION

Nematode parasites are nearly ubiquitous on commercial poultry farms. Surveys have found that 98.6% and 96% of commercial chicken farms are infected with the ascarid parasites *Ascaridia galli* and *Heterakis gallinarum*, respectively (Yazwinski et al., 2013). *A. galli* is a large (7-8 cm) ascarid nematode that lives in the small intestine, and although infections are typically subclinical, they can be associated with considerable production losses, impacting feed conversion, egg laying, as well as egg quality (Sharma et al., 2018; Collins et al., 2021). *H. gallinarum* is a small ascarid nematode (∼1 cm) that lives in the cecal pouches and causes no overt pathology or production losses in single species infections. However, *H. gallinarum* is a vector of the protozoan parasite *Histomonas meleagridis*, the causative agent of histomoniasis, more commonly known as blackhead disease (Tyzzer, 1934). Histomoniasis causes inflammation and necrosis of mucosal tissues and can be associated with significant mortality and production loss in infected animals, especially in turkeys (Tyzzer, 1934). To control the deleterious effects associated with *A. galli* and *H. gallinarum* infection, anthelmintics are used to treat nematode infections in poultry.

Currently, the only US Food and Drug Administration (FDA) approved treatment for nematode parasites in poultry is the benzimidazole (BZ) compound fenbendazole (FBZ) (Hoechst-Roussel-Vet, 2000; Hauck, 2024). Used for the past 25 years, FBZ has historically been associated with high efficacy against *A. galli, H. gallinarum*, as well as *Ascaridia dissimilis*, the large ascarid nematode of turkeys. However, resistance to BZ compounds is widespread in veterinary medicine and has previously been validated in ascarids of poultry (Crook et al., 2016; Collins et al., 2019, 2022). Loss of FBZ efficacy against *A. dissimilis* was first reported in 2013 and later validated in 2019 in a controlled efficacy study (Collins et al., 2019). In 2022, resistance to both the label and a double dose of FBZ was found in an *H. gallinarum* isolate (Collins et al., 2022). The emergence of poultry ascarid resistance to FBZ is of particular concern because of the lack of approved alternative treatments (Hauck, 2024). Without efficacious treatments, the detrimental effects associated with ascarid infections will proceed unabated, reducing the profit margins of poultry production. In addition, no treatments are available for histomoniasis, and FBZ-resistance in the *H. gallinarum* vector further limits control efforts. Given the importance of nematode management in poultry production, a broader survey of FBZ resistance on poultry farms is necessary to better understand the full scope of the problem.

Here, we have sampled thirteen isolates of *A. galli* and eight isolates of *H. gallinarum* from farms in South Carolina and Pennsylvania. We collected nematode embryos from each farm and tested FBZ efficacy in a controlled laboratory setting by infecting naïve animals and treating half of the animals with the label dosage of FBZ. Treatment efficacy was determined by counting nematode burdens in untreated and treated animals using the World Association for the Advancement of Veterinary Parasitology guidelines for poultry (Yazwinski et al., 2022). Resistance was detected in a single isolate of *A. galli* and all eight isolates of *H. gallinarum* tested, indicating that resistance is likely an emerging problem in *A. galli* but is highly prevalent in *H. gallinarum*.

## MATERIALS AND METHODS

### Collection of parasite isolates

Isolates of *A. galli* and *H. gallinarum* were collected from commercial broiler-breeder farms and diagnostic samples from layer and broiler-breeder farms, as well as one backyard flock (Fig 1, <u>S File 1</u>). Litter from farms or intestinal and cecal contents were first suspended in a 2:1 weight-to-volume ratio of tap water. The suspension was washed through two sieves, measuring 105 μM and 32 μM. The smaller of the sieves was monitored for lack of flow, at which point, the contents of the sieve were collected in 50 mL conical tubes. A plastic wash bottle filled with water was used to rinse the sieve. After rinsing, the remaining debris in the 105 μM sieve was discarded. The process was repeated until all of the suspension had been sieved. The 50 mL conicals were centrifuged at 1100 rpm (253 *g*) for three minutes to pellet the sample. Excess liquid was aspirated off the top of the pellet. A sodium nitrate solution (Vedco, FecaMed, St. Joseph, MO, specific gravity of ∼1.27) was then added to the pellet for a total tube volume of 45 mL. Samples were resuspended and then centrifuged at 1100 rpm (253 *g*) for three minutes to pellet the sample. The supernatant was poured over a 32 μM sieve, washed with water to remove sodium nitrate and then washed into a 15 mL collection tube using a bottle filled with distilled water. Embryos were allowed to settle, and then liquid was aspirated to leave a total volume of 5 mL of embryos in distilled water. Embryos were fully resuspended by vortexing, and five 20 μL aliquots were collected and placed on a microscope slide.

**Figure 1.**
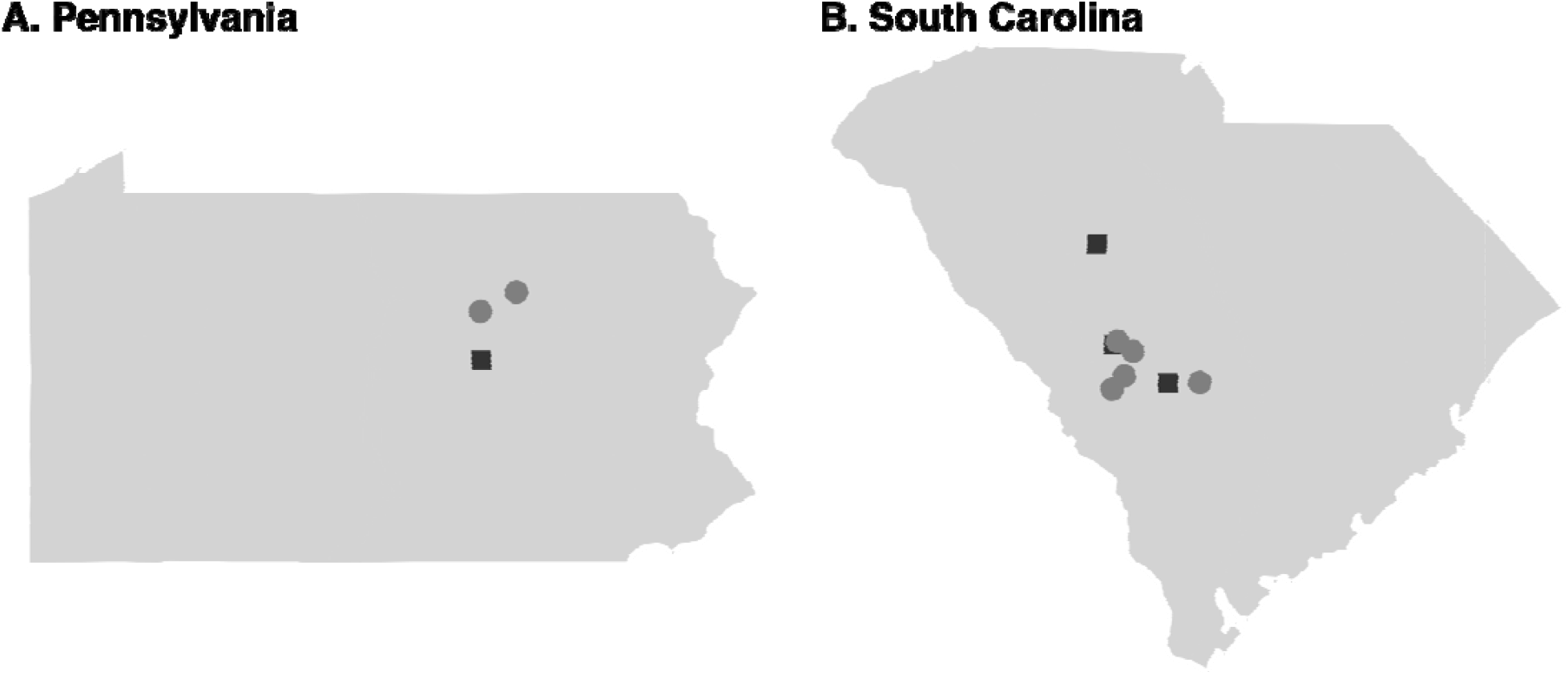
Location of farms sampled. Sampling locations are shown for A) Pennsylvania and B) South Carolina. Squares indicate that only *A. galli* was collected from the farm, and circles indicate that both *A. galli* and *H. gallinarum* were collected.

The number of embryos in each aliquot was counted, the average taken, and the estimated yield of embryos calculated. Embryos were resuspended in a total volume of 10 mL of distilled water and transferred to a 75 mL uncoated culture flask. Flasks were stored horizontally at room temperature for at least three weeks to allow embryo development to the infective stage with regular shaking and liquid added as needed.

### Infection and treatment of chickens

Animals were received as day-old chicks from either Sunnyside Hatchery (White leghorns) (Beaver Dam, WI) or Longenecker’s Hatchery (Ross 308) (Elizabethtown, PA) for each experiment and allowed to acclimate for one week. Animals were kept and handled under Johns Hopkins University IACUC approval (AV23A260). Each animal was then orally gavaged with ∼200 embryos of *A. galli* or *H. gallinarum* concentrated in 0.5 mL of water. Untreated and Treated animals were co-housed in the same room, and leg bands were used to distinguish infection and treatment groups. Animals were provided feed (Purina Unmedicated Start and Grow, St. Louis, MO) and water *ad libitum*. Four weeks after infection, animals in the groups to be treated were given FBZ by oral gavage (SafeGuard Aquasol, 1 mg/kg BW for five days) at a dosage representing 1.25 times the average body weight (BW) to account for variation and growth. Treatment was repeated at a similar time each day for five days.

### Necropsy and nematode quantification

Seven days after the final day of FBZ treatment, animals were humanely euthanized using carbon dioxide asphyxiation until a pulse was no longer detected. Cervical dislocation was used as a secondary method to confirm death. The intestine and ceca were collected from each animal by first making a medial cut into the skin under the keel. Skin and tissue were cut along each side of the breast, and then the breast was opened towards the head to open the body cavity. The fascia were cut, and the intestinal tract was pulled from the body cavity. The small intestine was collected by cutting a section from the duodenum to just below Meckel’s diverticulum. Ceca were removed by cutting at the ileocolic junction. The small intestine was opened longitudinally, and parasites were recovered and counted by gross examination of the contents. Cecal pouches were placed over a 32 μM sieve and sliced longitudinally. Cecal contents were rinsed using distilled water to remove contents from the tissue, and the contents were rinsed well through the sieve using distilled water. Nematodes collected on the sieve were washed into 50 mL conicals using distilled water, transferred to 10 cm petri dishes, and counted using a dissecting microscope.

### Statistical analysis

Efficacy of each isolate was determined as:

*Efficacy = ((Mean parasites per untreated bird)-(Mean parasites per treated bird))/(Mean Parasites per untreated bird)*

Mann-Whitney U-tests were performed in R (4.4.2) (R Core Team, 2020) to determine significant differences between the Untreated and Treated groups for each isolate.

## RESULTS

### Ascaridia galli

Isolates of *A. galli* were collected from thirteen farms in either South Carolina or Pennsylvania. After five days of FBZ treatment, infected animals were humanely euthanized, and parasite burdens were quantified. Efficacy was calculated, and statistical comparisons between the Untreated and Treated groups for each isolate were made using the Mann-Whitney U-test. Twelve isolates were found to be 100% FBZ susceptible, with the caveat that SC7 had low parasite establishment in the Untreated group. However, the C+E isolate was found to have significantly reduced efficacy (67%) (Table 1, Fig 2), and no significant differences were observed between the Untreated and Treated groups (*p* = 0.49).

**Table 1.** Efficacy of FBZ against each *A. galli* isolate. Resistant isolates are shown in red. Standard error (SE) shown in parentheses.

| Farm | Avg. parasites<br>Untreated (SE) | Avg. parasites Treated<br>(SE) | Efficacy (SE) |
| --- | --- | --- | --- |
| SC1 | 4 (0.71) | 0 (0) | 100% (25) |
| SC2 | 4.83 (1.62) | 0 (0) | 100% (47.43) |
| SC3 | 33.83 (10.87) | 11.5 (6.66) | 66% (43.24) |
| SC4 | 10.14 (3.2) | 0 (0) | 100% (44.65) |
| SC5 | 35.6 (8.52) | 0 (0) | 100% (33.84) |
| SC6 | 13.4 (2.66) | 0 (0) | 100% (28.04) |
| SC7 | 0.14 (0.14) | 0 (0) | 100% (141.42) |
| SC8 | 9.4 (1.86) | 0 (0) | 100% (27.98) |
| SC9 | 9.17 (4.56) | 0 (0) | 100% (70.41) |
| SC10 | 16.75 (5.24) | 0 (0) | 100% (45.12) |
| PA1 | 21.8 (11.53) | 0 (0) | 100% (74.77) |
| PA2 | 9.75 (3.14) | 0.14 (0.14) | 98.53% (45.29) |
| PA3 | 7.6 (1.83) | 0 (0) | 100% (34.11) |

**Figure 2.**
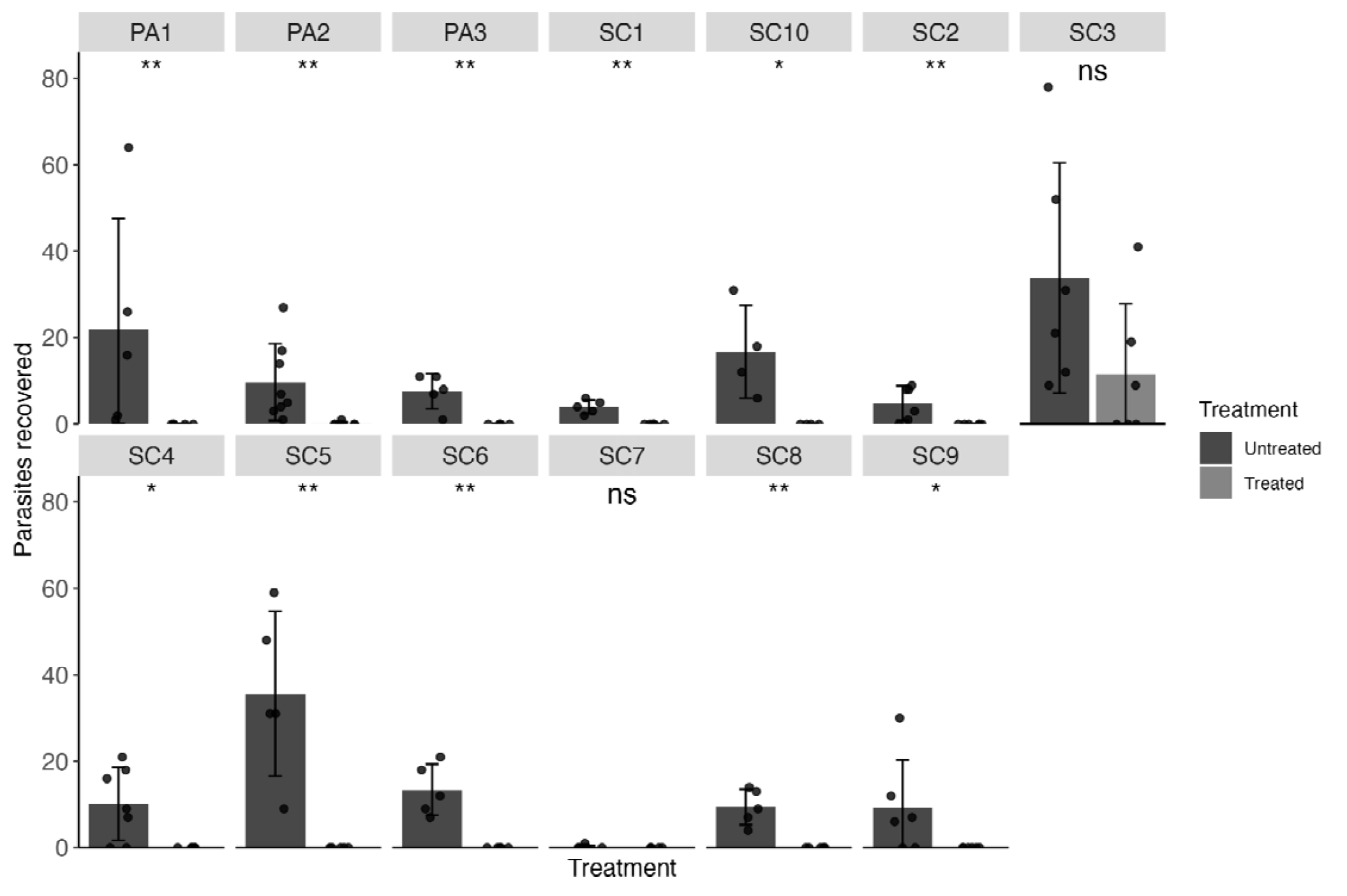
FBZ efficacy (1.25 mg/kg body weight over five days) in isolates of *A. galli*, faceted by farm of origin. Means are shown as bars for each farm with standard deviations. The number of parasites recovered from each infected animal is shown as points. Statistical significance is shown above each treatment comparison (*p* > 0.05 = ns, *p* < 0.05 = *, *p* < 0.01 = **, Mann-Whitney U test).

### H. gallinarum

Isolates of *H. gallinarum* were collected from thirteen farms in either South Carolina or Pennsylvania. Evaluation and analysis of FBZ efficacy was performed as for *A. galli*. All eight isolates screened were found to be resistant to FBZ treatment, with efficacies ranging from 19.35 to 66% (Table 2, Fig 3).

**Table 2.**
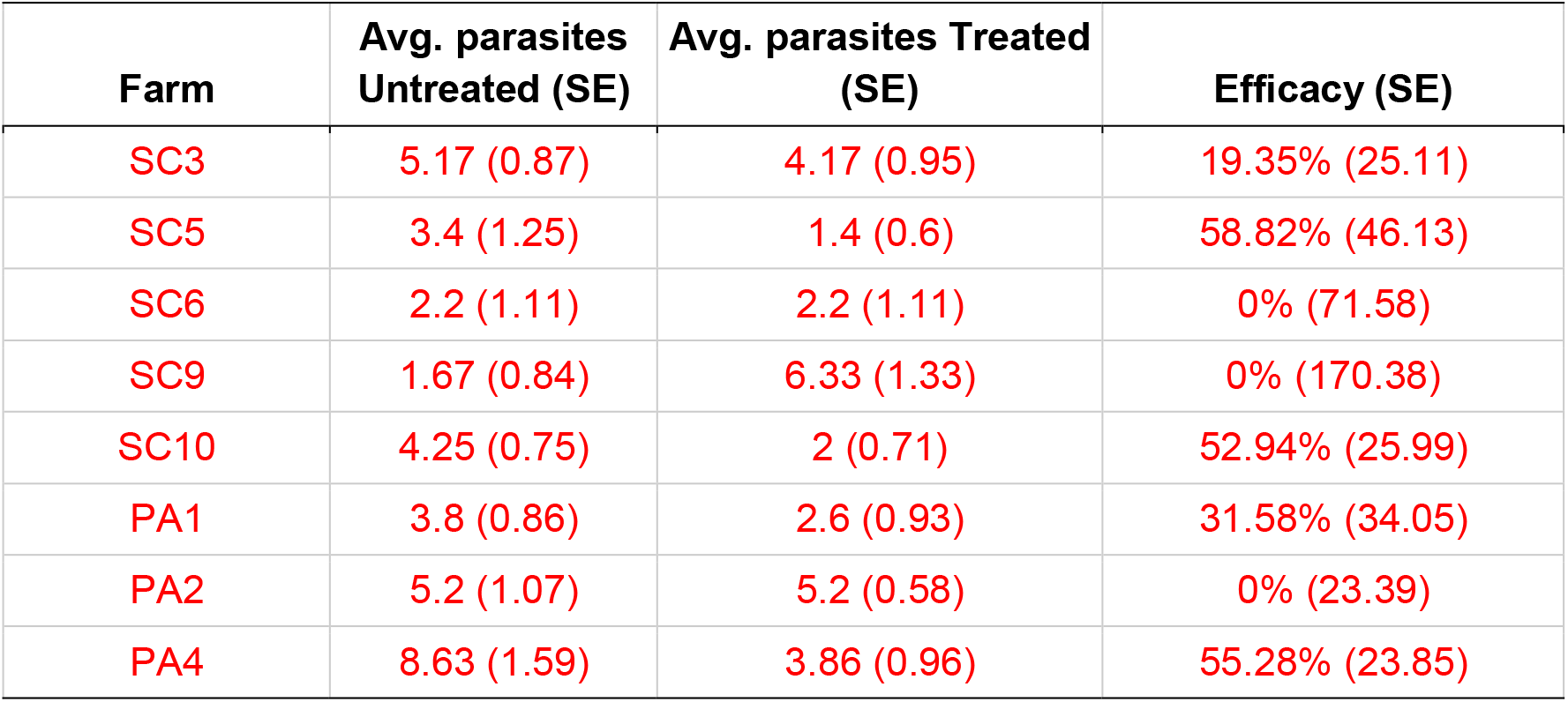
Efficacy of FBZ against each *H. gallinarum* isolate. Resistant isolates are shown in red. Standard error (SE) shown in parentheses.

| Farm | Avg. parasites Untreated (SE) | Avg. parasites Treated (SE) | Efficacy (SE) |
| --- | --- | --- | --- |
| SC3 | 5.17 (0.87) | 4.17 (0.95) | 19.35% (25.11) |
| SC5 | 3.4 (1.25) | 1.4 (0.6) | 58.82% (46.13) |
| SC6 | 2.2 (1.11) | 2.2 (1.11) | 0% (71.58) |
| SC9 | 1.67 (0.84) | 6.33 (1.33) | 0% (170.38) |
| SC10 | 4.25 (0.75) | 2 (0.71) | 52.94% (25.99) |
| PA1 | 3.8 (0.86) | 2.6 (0.93) | 31.58% (34.05) |
| PA2 | 5.2 (1.07) | 5.2 (0.58) | 0% (23.39) |
| PA4 | 8.63 (1.59) | 3.86 (0.96) | 55.28% (23.85) |

**Figure 3.**
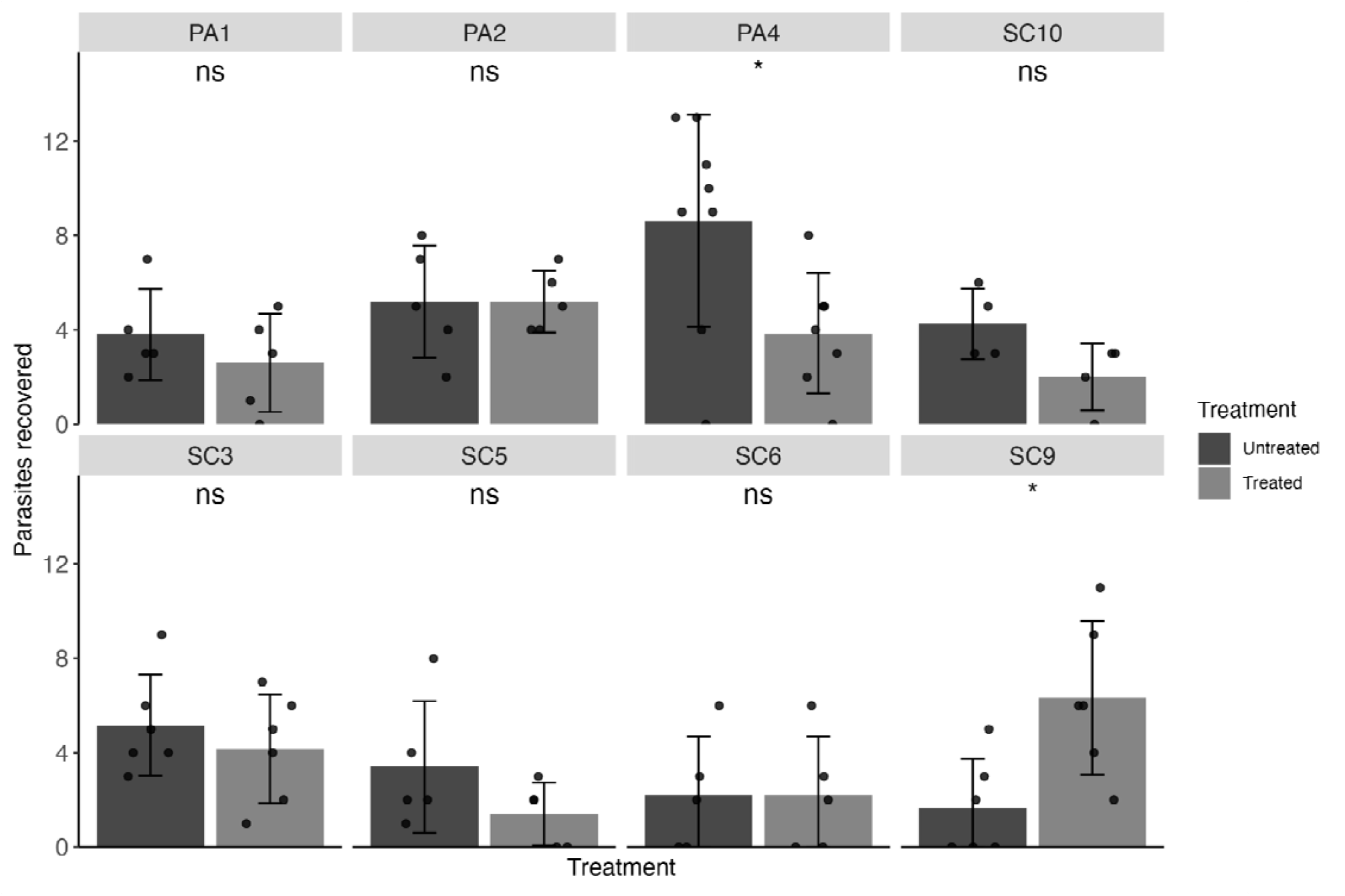
Fenbendazole efficacy (1.25 mg/kg body weight over five days) in isolates of *Heterakis gallinarum*, faceted by farm of origin. Means are shown as bars for each farm with standard deviations. The number of parasites recovered from each infected animal is shown as points. Statistical significance is shown above each treatment comparison (*p* > 0.05 = ns, *p* < 0.05 = *, *p* < 0.01 = **, Mann-Whitney U test).

## DISCUSSION

Despite the near ubiquity of BZ resistance in many nematodes of veterinary importance, BZ resistance in ascarids remains poorly studied. We have previously confirmed and validated FBZ resistance in *A. dissimilis* and *H. gallinarum* (Collins et al., 2019, 2022). Here, we go one step further to examine efficacy across several farms, documenting the first confirmed case of resistance in *A. galli* and highlighting widespread resistance in *H. gallinarum*.

We screened thirteen isolates of *A. galli* and eight isolates of *H. gallinarum* to determine FBZ efficacy in a controlled research setting. We identified a single resistant isolate of *A. galli*, indicating that resistance is likely an emerging problem and prevalence is likely to increase without new treatments. Additionally, *A. galli* is the third ascarid species of poultry with a validated resistant isolate. Resistance in *A. galli* and *H. gallinarum* has so far been found only in isolates collected from either commercial layers or broiler-breeders (Collins et al., 2022). Similar to commercial turkey production, where FBZ-resistant *A. dissimilis* was found, both production types involve birds living for an extended periods compared to commercial broilers. Each time a parasite population is exposed to treatment, selection is applied and mutations that confer resistance increase in frequency over time. The extended lifespans in breeders and layers necessitate repeated treatments of the same birds and parasite populations, which amplifies selective pressure, increasing the odds of resistance emerging (Jackson and Coop 2000). Over time, resistance is likely to emerge in parasites on broiler farms, albeit likely more slowly because of reduced selective pressures.

Although not a major health concern in poultry, the emergence of resistance in *A. galli* could have a substantial impact on production because of the potential economic costs associated with FBZ resistance. In *A. dissimilis-*infected turkeys, when feed conversion was compared between animals infected with either a susceptible or resistant isolate, animals infected with resistant parasites and treated on a typical schedule had significantly decreased feed conversion over a 10-week growth period (Collins et al., 2021). If new interventions are not found, chicken production could face diminishing profit margins.

All eight isolates of *H. gallinarum* screened were found to be resistant, demonstrating potential differences in how parasite burdens accumulate and persist in poultry. *H. gallinarum* burdens have been shown to persist throughout the bird’s lifetime (Stehr et al., 2018), and in longer-lived animals, selection on parasites is likely stronger due to repeated use of FBZ throughout the host’s life. Resistance in *H. gallinarum* is of particular concern because of its role as a vector for *H. meleagridis*. Although transmission of histomoniasis within a flock is typically from bird to bird, *H. meleagridis* quickly dies in the environment (Lotfi et al., 2012), making transmission of *H. meleagridis* infections to subsequent flocks dependent on protozoa surviving in the embryos of the ascarid vector. The removal of effective arsenical treatments for histomoniasis is believed to have led to an increase in prevalence in chickens (Grafl et al., 2011; Dolka et al., 2015; Clark and Kimminau, 2017). Our results indicate that loss of effective vector control is likely an underlying cause of increased prevalence of histomoniasis.

Overall, we have demonstrated that resistance is present across all three species of poultry ascarids and, within our small sampling of populations, is ubiquitous in *H. gallinarum*. However, these data represent a few farms within two states and are not a representation of the industry at large. Large-scale sampling across all production types throughout the US is necessary to ascertain the full scope of resistance. Broad-scale *in vivo* screenings, such as those performed here, of hundreds of farms are not feasible because of the costs associated with performing controlled efficacy testing, necessitating a higher-throughput method of screening for resistance. In parasites of small ruminants, high-throughput diagnostics for resistance have been created using deep-amplicon sequencing (Avramenko et al., 2019), which enables sequence-based analysis for known resistance markers for many isolates at one time. However, our preliminary data and published reports from the horse ascarid *Parascaris univalens* (Martin et al., 2021) indicate that FBZ-resistance mechanisms are unique in ascarids when compared to *H. contortus*. Therefore, it is necessary to study the genetics of ascarids to determine mechanisms of BZ resistance so that diagnostic tools can be developed using a defined set of resistance markers in ascarids. Using new diagnostics, broad surveys of different poultry production types can be conducted to obtain more significant sampling of the prevalence of BZ resistance in poultry ascarids. Insights from such surveys could then be used to develop new management strategies for FBZ-resistant parasites, as well as act as an impetus for the development and use of new treatments in poultry production.

## ACKNOWLEDGMENTS

We would like to thank the US Poultry and Egg Association to ECA, National Institute of Health (R21 AI180805 to ECA), and United States Department of Agriculture (2024-67012-43769 to JBC) for funding. Additionally, we thank Amick Farms and the Pennsylvania State Animal Diagnostic Lab for facilitating the collection of screened isolates.

## REFERENCES

Avramenko, R. W., E. M. Redman, L. Melville, Y. Bartley, J. Wit, C. Queiroz, D. J. Bartley, and J. S. Gilleard. 2019. Deep amplicon sequencing as a powerful new tool to screen for sequence polymorphisms associated with anthelmintic resistance in parasitic nematode populations. Int. J. Parasitol. 49:13–26 Available at http://dx.doi.org/10.1016/j.ijpara.2018.10.005.

Clark, S., and E. Kimminau. 2017. Critical Review: Future Control of Blackhead Disease (Histomoniasis) in Poultry. Avian Dis. 61:281–288 Available at http://dx.doi.org/10.1637/11593-012517-ReviewR.

Collins, J. B., B. Jordan, L. Baldwin, C. Hebron, K. Paras, A. N. Vidyashankar, and R. M. Kaplan. 2019. Resistance to fenbendazole in Ascaridia dissimilis, an important nematode parasite of turkeys. Poult. Sci. 98:5412–5415 Available at https://www.ncbi.nlm.nih.gov/pubmed/31328783.

Collins, J. B., B. Jordan, A. Vidyashankar, A. Bishop, and R. M. Kaplan. 2022. Fenbendazole resistance in Heterakis gallinarum, the vector of Histomonas meleagridis, on a broiler breeder farm in South Carolina. Vet Parasitol Reg Stud Reports 36:100785 Available at http://dx.doi.org/10.1016/j.vprsr.2022.100785.

Collins, J. B., B. Jordan, A. N. Vidyashankar, P. J. Castro, J. Fowler, and R. M. Kaplan. 2021. Impact of fenbendazole resistance in Ascaridia dissimilis on the economics of production in turkeys. Poult. Sci. 100:101435 Available at http://dx.doi.org/10.1016/j.psj.2021.101435.

Crook, E. K., D. J. O’Brien, S. B. Howell, B. E. Storey, N. C. Whitley, J. M. Burke, and R. M. Kaplan. 2016. Prevalence of anthelmintic resistance on sheep and goat farms in the mid-Atlantic region and comparison of in vivo and in vitro detection methods. Small Rumin. Res. 143:89–96 Available at https://www.sciencedirect.com/science/article/pii/S0921448816302413.

Dolka, B., A. Zbikowski, I. Dolka, and P. Szeleszczuk. 2015. Histomonosis - an existing problem in chicken flocks in Poland. Vet. Res. Commun. 39:189–195 Available at https://www.ncbi.nlm.nih.gov/pubmed/25976057.

Grafl, B., D. Liebhart, M. Windisch, C. Ibesich, and M. Hess. 2011. Seroprevalence of Histomonas meleagridis in pullets and laying hens determined by ELISA. Vet. Rec. 168:160 Available at http://dx.doi.org/10.1136/vr.c6479.

Hauck, R. 2024. Helminthiasis in Poultry. Merck Veterinary Manual Available at https://www.merckvetmanual.com/poultry/helminthiasis/helminthiasis-in-poultry (verified 15 July 2025).

Hoechst-Roussel-Vet. 2000. FREEDOM OF INFORMATION SUMMARY, SUPPLEMENTAL NEW ANIMAL DRUG APPLICATION, NADA 131-675, SAFE-GUARDÂ® (Fenbendazole) For Growing Turkeys.

Lotfi, A.-R., E. M. Abdelwhab, and H. M. Hafez. 2012. Persistence of Histomonas meleagridis in or on materials used in poultry houses. Avian Dis. 56:224–226 Available at http://dx.doi.org/10.1637/9519-090910-ResNote.1.

Martin, F., P. Halvarsson, N. Delhomme, J. Höglund, and E. Tydén. 2021. Exploring the β-tubulin gene family in a benzimidazole-resistant Parascaris univalens population. Int. J. Parasitol. Drugs Drug Resist. 17:84–91 Available at http://dx.doi.org/10.1016/j.ijpddr.2021.08.004.

R Core Team. 2020. R: A Language and Environment for Statistical Computing. Available at https://www.R-project.org/.

Sharma, N., P. W. Hunt, B. C. Hine, N. K. Sharma, A. Chung, R. A. Swick, and I. Ruhnke. 2018. Performance, egg quality, and liver lipid reserves of free-range laying hens naturally infected with Ascaridia galli. Poult. Sci. 97:1914–1921 Available at https://www.ncbi.nlm.nih.gov/pubmed/29562346.

Stehr, M., Q. Sciascia, C. C. Metges, M. Gauly, and G. Das. 2018. Co-expulsion of Ascaridia galli and Heterakis gallinarum by chickens. Int. J. Parasitol. 48:1003–1016 Available at https://www.ncbi.nlm.nih.gov/pubmed/30240707.

Tyzzer, E. E. 1934. Studies on Histomoniasis, or “Blackhead” Infection in the Chicken and the Turkey. Proceedings of the American Academy of Arts and Sciences 69:189–264 Available at http://www.jstor.org/stable/20023041 (verified 28 January 2020).

Yazwinski, T. A., J. Höglund, A. Permin, M. Gauly, and C. Tucker. 2022. World association for the advancement of veterinary parasitology (WAAVP): Second edition of guidelines for evaluating the efficacy of anthelmintics in poultry. Vet. Parasitol. 305:109711 Available at http://dx.doi.org/10.1016/j.vetpar.2022.109711.

Yazwinski, T., C. Tucker, E. Wray, L. Jones, Z. Johnson, S. Steinlage, and J. Bridges. 2013. A survey on the incidence and magnitude of intestinal helminthiasis in broiler breeders originating from the southeastern United States1. J. Appl. Poult. Res. 22:942–947 Available at http://dx.doi.org/10.3382/japr.2013-00776.

